# Documenting coral spawning in East Africa: New *in situ* observations from Zanzibar for three reef-building species

**DOI:** 10.1101/2025.10.02.679943

**Authors:** Ashlee Lillis, Narriman Jiddawi

## Abstract

Broadcast coral spawning is a vital reproductive event for many reef-building species and is essential to the resilience of coral reef ecosystems. Understanding spawning dynamics is key to assessing the reproductive health and resilience of these ecosystems. While extensively documented in the Indo-Pacific and Caribbean, coral spawning remains understudied in the Western Indian Ocean (WIO), a region facing rapid environmental change and increasing coral decline. This study presents the first *in situ* coral spawning records from Zanzibar for three Scleractinian species: *Galaxea astreata, Favites pentagona*, and *Platygyra daedalea*. Observations were made during monthly nighttime monitoring dives from November 2024 to February 2025, timed with lunar and solar cycles linked to peak spawning in other regions. Results showed species-specific spawning timing: *G. astreata* spawned on the fourth night after the full moon in November and December; *F. pentagona* on the sixth night in November; and *P. daedalea* on the sixth night in December and January. Spawning behaviour, including gamete setting, release windows, and split spawning, also varied by species. These findings offer the first detailed insights into reproductive patterns for these species in the WIO and underscore the importance of local spawning studies to support regional coral conservation and restoration.

## 1. Introduction

Coral reef ecosystems are among the most biodiverse on Earth and provide innumerable ecological services to coastal and island populations, from fish nursery habitat to shoreline protection (Bellwood *et al*., 2004; Burke and Spalding, 2022; Santavy *et al*., 2021). Coastal ecosystems, and coral reefs in particular, are in rapid decline due to myriad stressors both at local (e.g., coastal development, run-off, sedimentation, water quality, destructive fishing) and regional or global scales (e.g., climate change, invasive species). Coral reef degradation is predicted to continue unless these threats can be managed and recovery can be facilitated (Bellwood *et al*., 2004). The foundation of these critical ecosystems are reef-building stony corals, sessile invertebrate animals that live in symbiosis with photosynthetic organisms called zooxanthellae and produce a calcium carbonate skeleton.

East African coral reefs are an important yet understudied sub-region for coral diversity and connectivity (Mbije *et al*., 2002). In Zanzibar coral reefs are also a highly valuable coastal resource, increasingly experiencing widespread degradation due to various human stressors (Ussi *et al*., 2024). Though no comprehensive census has been completed for Zanzibar, there are currently estimated between 150-175 hard coral species (Johnstone *et al*., 1998; Khamis *et al*., 2017) and despite declines over the past 30 years healthy coral formations persist, especially in more remote areas protected from human impacts (Bravo *et al*., 2021; Muhando, 2009; Ussi *et al*., 2024). Key to continued persistence and the resilience potential of reefs is successful coral reproduction. Most of the reef-building hard coral species are broadcast spawners, simultaneously releasing eggs and sperm into the water column typically only once or twice per year, where the eggs are fertilized in surface waters and develop through a larval stage before settling and attaching to the seafloor (Babcock *et al*., 1986; Baird *et al*., 2021; Harrison and Wallace, 1990). These synchronized spawning events are known to occur across vast expanses of reef. Where dense coral populations exist, large surface slicks of coral gametes can be observed at the water surface following a spawning event (Harrison *et al*., 1984; Jamodiong *et al*., 2018).

Spawning is closely synchronized to solar, lunar, and seasonal cycles, and under the influence of environmental factors such as water temperature and light (Baird *et al*., 2009; Boch *et al*., 2011; Harrison *et al*., 1984; Lin and Nozawa, 2017, 2023; Penland et al., 2004). For example, a given species may consistently spawn annually on the 10^th^ night after the full moon in May between 45-60 minutes after sunset (Baird *et al*., 2009; Banaszak *et al*., 2023). Because gametes only remain viable for a few hours, it is critical to reproductive success that individuals in a coral species spawn in such close coordination. In the most well studied regions (e.g., the Great Barrier Reef), mass spawning of dozens of species has been observed over just one or two nights in a single year (Baird *et al*., 2021; Harrison *et al*., 1984); however, variation in spawning frequency and timing, as well as inter-annual consistency, is known to occur across geographies (Lin and Nozawa, 2017, 2023; Sakai *et al*., 2020) and there are instances of populations with protracted non-overlapping spawning periods of several months (Mangubhai *et al*., 2007; Mangubhai and Harrison, 2008, 2009).

Understanding coral reproductive phenology and spawning behaviour has become imperative in recent decades as scientists and conservation practitioners have increased efforts to stem declines in coral populations and to regenerate reef habitat through active restoration (Banaszak *et al*., 2023; Miller *et al*., 2024). Traditional restoration practices rely upon asexual propagation, cloning existing coral via fragmentation, which does not directly increase genetic diversity or create genetically new individuals. Because sexual reproduction can produce hundreds of millions of coral propagules at a single time from many parents, it provides new genetic diversity that promotes the population resilience required to replenish and sustain future reef communities (Banaszak *et al*., 2023). Restoration projects have expanded to include sexual propagation in many areas of the Caribbean and the Indo-Pacific (Banaszak *et al*., 2023; Miller *et al*., 2022), and sexually propagated recruits have recently shown higher thermal tolerance during bleaching events compared to adult and asexually propagated coral (Miller *et al*., 2024). Coral restoration efforts in the Western Indian Ocean, including Zanzibar and mainland Tanzania, are increasing rapidly; however, all currently known projects use only asexual propagation via fragmentation.

Coral reproductive patterns in East Africa remain largely undocumented, despite the ecological importance of its biodiverse nearshore reefs. In Zanzibar, establishing baseline data on spawning timing is critical for understanding the reproductive potential of local coral populations and informing restoration strategies that incorporate sexual propagation—a key element for enhancing reef resilience (Banaszak *et al*., 2023). Although over 6,500 coral spawning records exist across the Indo-Pacific (Baird *et al*., 2021), empirical data from the Western Indian Ocean remain extremely limited. To date, only two opportunistically observed spawning events have been reported from Zanzibar, and no direct *in situ* observations have been published for East Africa. The only prior studies in the region inferred spawning timing indirectly through histological analyses of gametogenesis and post-settlement monitoring of coral spat in Kenya’s Mombasa Marine Reserve (Mangubhai *et al*., 2007; Mangubhai and Harrison, 2008b). To address this significant knowledge gap, we conducted the first targeted *in situ* observations of coral spawning in Zanzibar, documenting the timing and gamete release behaviours of three Scleractinian species for which no prior spawning records exist in the Western Indian Ocean.

## 2. Materials and Methods

### Study Area: Jambiani, Zanzibar, Tanzania

Zanzibar is a semi-autonomous region of the United Republic of Tanzania, comprising two major tropical islands (Unguja and Pemba) and many small ones (Fig. 1a). The Zanzibar archipelago has substantial fringing and patch coral reefs, estimated to cover 90 km^2^, hosting high biodiversity and providing significant economic value to humans (Bravo *et al*., 2021; Khamis *et al*., 2017). The reef systems are key to support local fisheries, tourism, and provide coastal protection (Bravo *et al*., 2021; Khamis *et al*., 2017). The eastern coasts of the two major islands have continuous reefs fringing the coastline and creating protected shallow lagoons with wide beaches.

**Figure 1.**
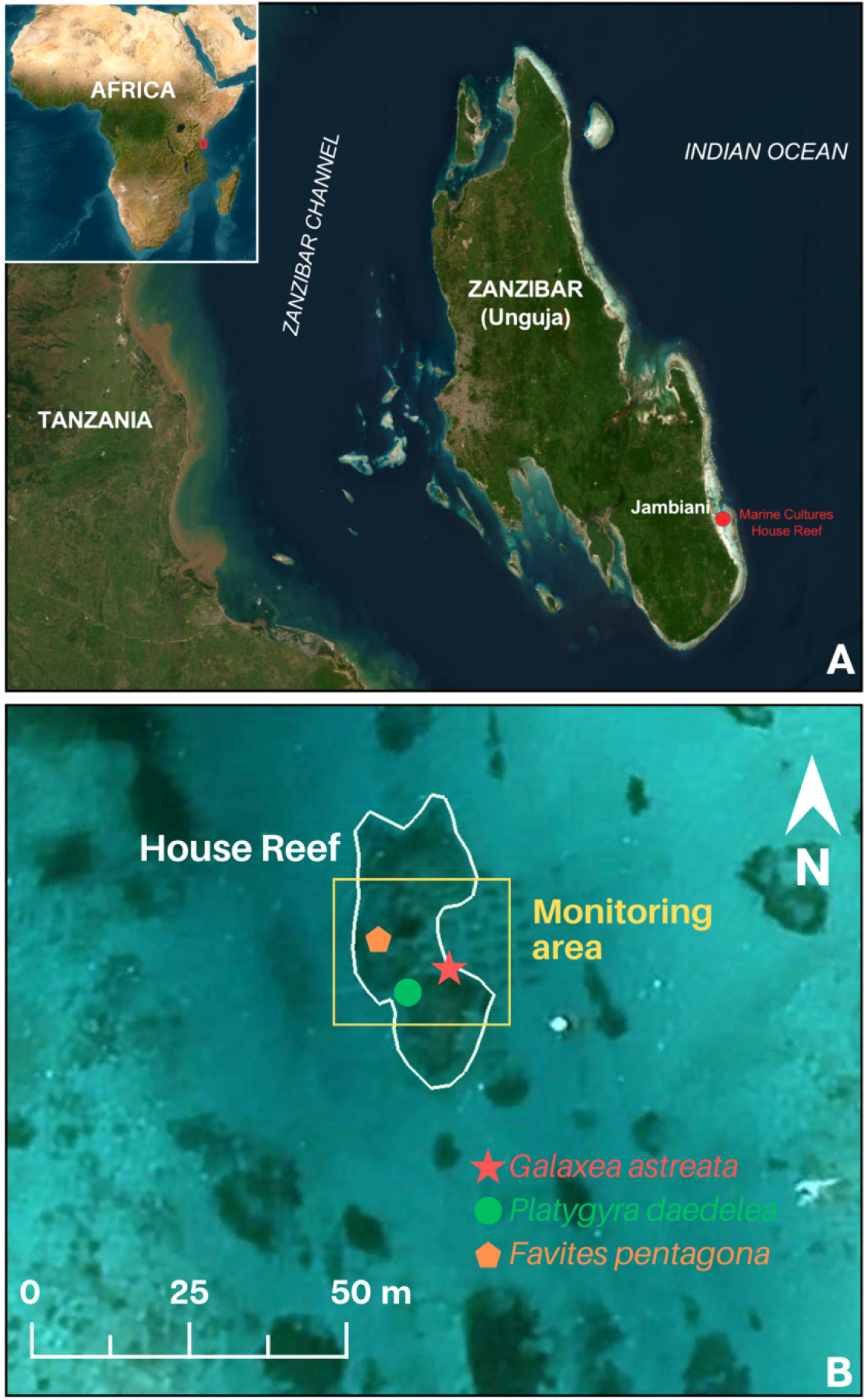
A. Map of Unguja Island, Zanzibar showing location of Jambiani village and study reef site (red). B. Satellite image of the Marine Cultures House Reef with monitoring area delineated. Locations of each coral species observed spawning are indicated.

This study was conducted at a shallow patch reef site in Jambiani lagoon. Jambiani, a coastal village bordering a shallow lagoon, is located on the southeast coast of Unguja Island (Fig.1a). The lagoon is formed by a fringing coral reef system several kilometers offshore creating a barrier from the open ocean, while the lagoon comprises patches of coral reef, seagrass, and soft bottom. Traditionally a fishing village, the area has seen rapid coastal development and environmental change in the past 10-15 years. This lagoon system experiences large tidal fluctuations (2-4 m tidal range), seasonal sedimentation, and is hydrodynamically active.

Zanzibar-based marinecultures.org is a non-profit organization operating a coral farm and transplantation program in Jambiani lagoon (Fig.1). Beginning in 2014 corals were fragmented, grown on nursery tables, and planted to their “house reef” (MC reef; Fig.1b). The MC reef is approximately 750 m^2^ (50 m x 15 m), ranging from 2-9 m depth, with established large (>5 m diameter) colonies of *Porites* spp., *Galaxea astreata*, and *Lobophyllia* sp., and many smaller (0.5-1 m diameter) colonies of a variety of species.

This patch reef, while relatively small, is estimated to contain individuals of ∼40 species of hard coral, including at least 30 genera. Like other reefs around the world, Zanzibar recently experienced a mass coral bleaching event during 2024, severely impacting the MC reef site where the majority of planted Acroporids perished. The coral cover is now estimated at 35-40%, *Acropora* spp. are rare, and many surviving coral colonies show partial mortality.

### Field Observations

A series of monthly night dives on SCUBA, beginning just after sunset, were carried out during November and December 2024, January and February 2025 at the MC site, totaling 1166 minutes of observation. Dives were scheduled each month for the 3-6 nights after the full moon (NAFM), based on the highest number of observations of coral spawning during these nights in other locales (Baird et al., 2021, 2022). Lunar phases and local timing for Zanzibar were obtained from timeanddate.com. On each observation night, two divers descended at sunset to the reef site, each patrolling a small area (∼50m^2^) for the duration of the dive (1-2 hours depending on sea conditions and air consumption) to consistently monitor the same coral colonies each dive. Divers monitored corals within their zone to check for “setting”, when gamete bundles appear in polyp mouths prior to release, as well as “spawning”, when positively buoyant spherical gamete bundles are released into the water column and float towards the surface. While coral spawning can be obscured by high amounts of turbidity and light-attracted plankton in the water, gamete bundles observed were distinct and were confirmed to be originating from coral.

Time-stamped photos and video were taken to document any setting or spawning behaviours observed, as well as the conclusion of any spawning event. Observations included data on setting and spawning timing, duration, and synchrony (when multiple individuals of the same species were present). The same areas, and therefore same coral colonies, were observed each month of observation.

This work was carried out under Zanzibar Research Permit *#*2001709162242701335980.

## 3. Results

Three hermaphroditic coral species were recorded spawning during this initial monitoring interval at MC reef, each with distinct timing and release behaviour (Table 1). The repeated monitoring of the same individual corals enabled detailed observations of their specific gamete release patterns. Spawning events for each species observed are described in detail in individual sections below.

**Table 1.** Summary data for coral spawning. N = number of coral colonies observed spawning, * indicates that the same colony was observed spawning on more than one occasion. NAFM = nights after full moon. MAS = minutes after sunset.

| Species | Date(s) of Observed Spawn | N | NAFM | Time of spawn (start) | MAS | Spawning period (min) | Gamete setting observed? |
| --- | --- | --- | --- | --- | --- | --- | --- |
| *Galaxea astreata* | 19-Nov-2024 | 1* | 4 | 19:07 | 46 | 31 | N |
|  | 19-Dec-2024 | 1* | 4 | 19:15 | 54 | 20 | N |
| *Favites pentagona* | 21-Nov-2024 | 1 | 6 | 19:24 | 63 | 11 | Y (19:13) |
| *Platygyra daedalea* | 21-Dec-2024 | 1* | 6 | 19:56 | 80 | 3 | N |
|  | 21-Dec-2024 | 1* | 6 | 20:26 | 110 | 2 | Y (20:00) |
|  | 20-Jan-2025 | 4* | 6 | 20:06<br>20:11<br>20:19 | 80 | 2 | Y (19:28) |

### Species 1: *Galaxea astreata* (Family Euphylliidae)

A large (∼8m diameter) colony of *G. astreata* was observed spawning on the 4^th^ nights after the full moon in November and December 2024 (Table 1, Fig.2). This colony had experienced severe bleaching in April 2024, leading to approximately 30-40% mortality. Spawning was detected 46 and 54 minutes after sunset and continued for 31 and 20 minutes in November and December, respectively. Gamete bundles were white in colour and seen rising from the large colony. Due to the coral’s morphology and long extended tentacles, it was not possible to see gamete bundles emerge from polyp mouths in this species. The December spawning event was observed to be smaller (i.e., fewer gamete bundles released) with gamete bundles appearing to be smaller in diameter than during the larger spawn in November.

**Figure 2.**
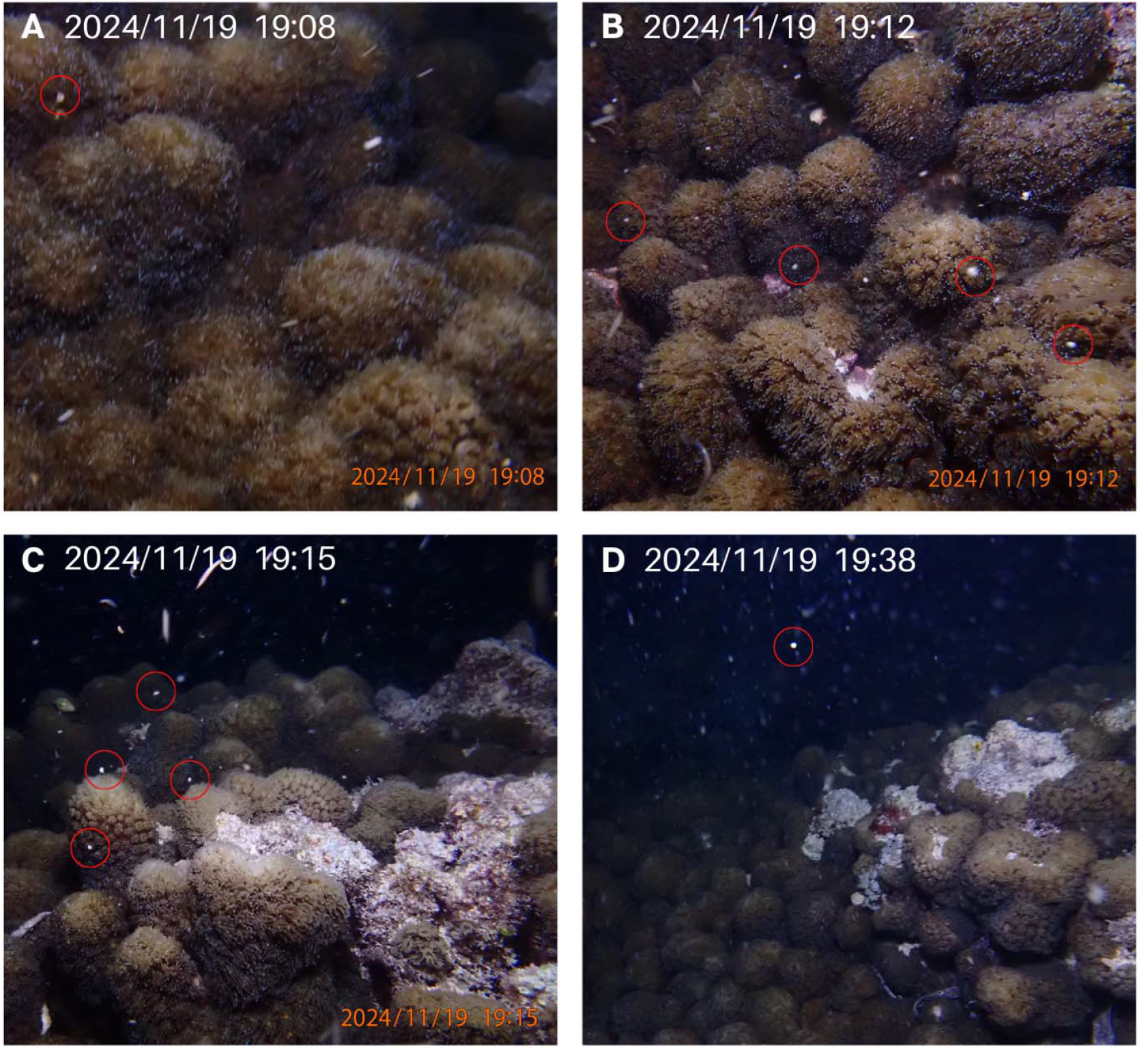
*Galaxea astreata* November 2024 spawning event. White coloured gamete bundles rising in the water column are circled in red. Recent mortality of the colony can be seen in C and D.

### Species 2: *Favites pentagona* (Family Merulinidae)

On the 6^th^ night after the full moon in November 2024, ripe pink gamete bundles were observed setting (visible in polyp mouths) at 19:13 in a single colony of *F. pentagona* (Fig. 3A). Divers searched the monitoring area for additional colonies of *F. pentagona* but none were found, therefore it was only possible to observe one individual. First gamete release began at 19:24 (Fig. 3C), 14 minutes after setting was noted, and continued until the final bundle was released at 19:35.

**Figure 3.**
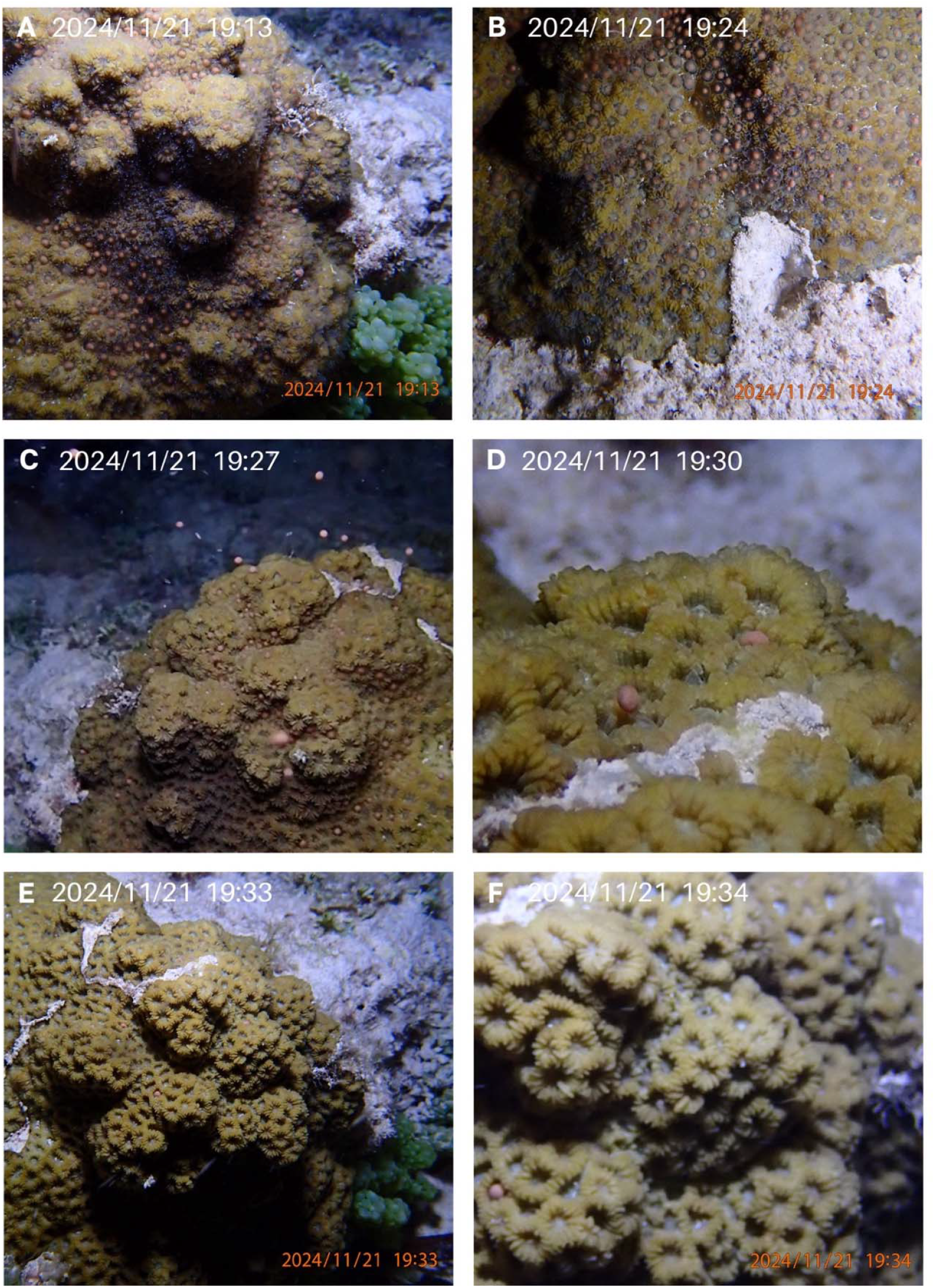
*Favites pentagona* spawning sequence in November 2024. Gamete setting is seen in panel A. First gamete bundles were released at 19:24 (B) and continued for 11 minutes (C-E), with the final bundle released seen in F.

To check if this *F. pentagona* colony would spawn again at the same lunar timing in December, it was monitored 3-6 nights after the full moon from 30-90 minutes after sunset. No spawning was observed.

### Species 3: *Platygyra daedalea* (Family Merulinidae)

A brain coral species common to the MC reef, *P. daedalea*, spawned in December 2024 and in January 2025, on the 6^th^ night after the full moon, starting at 80 minutes after sunset in both months (Table 1, Fig. 4,5). Gamete release was remarkably rapid in this species, with pulses of spawning lasting only 2-3 minutes. Five adjacent colonies of this species are present in a small area (< 5 m^2^) and were able to be monitored simultaneously. In December, a single individual colony was observed spawning. Gamete release had already begun when the diver came across the spawning colony, thus initial setting was not observed. However, only part of the colony spawned in the first pulse at 80 minutes after sunset, while other parts of the colony began setting shortly after with spawning occurring again 30 minutes later (Table 1, Fig. 4).

**Figure 4.**
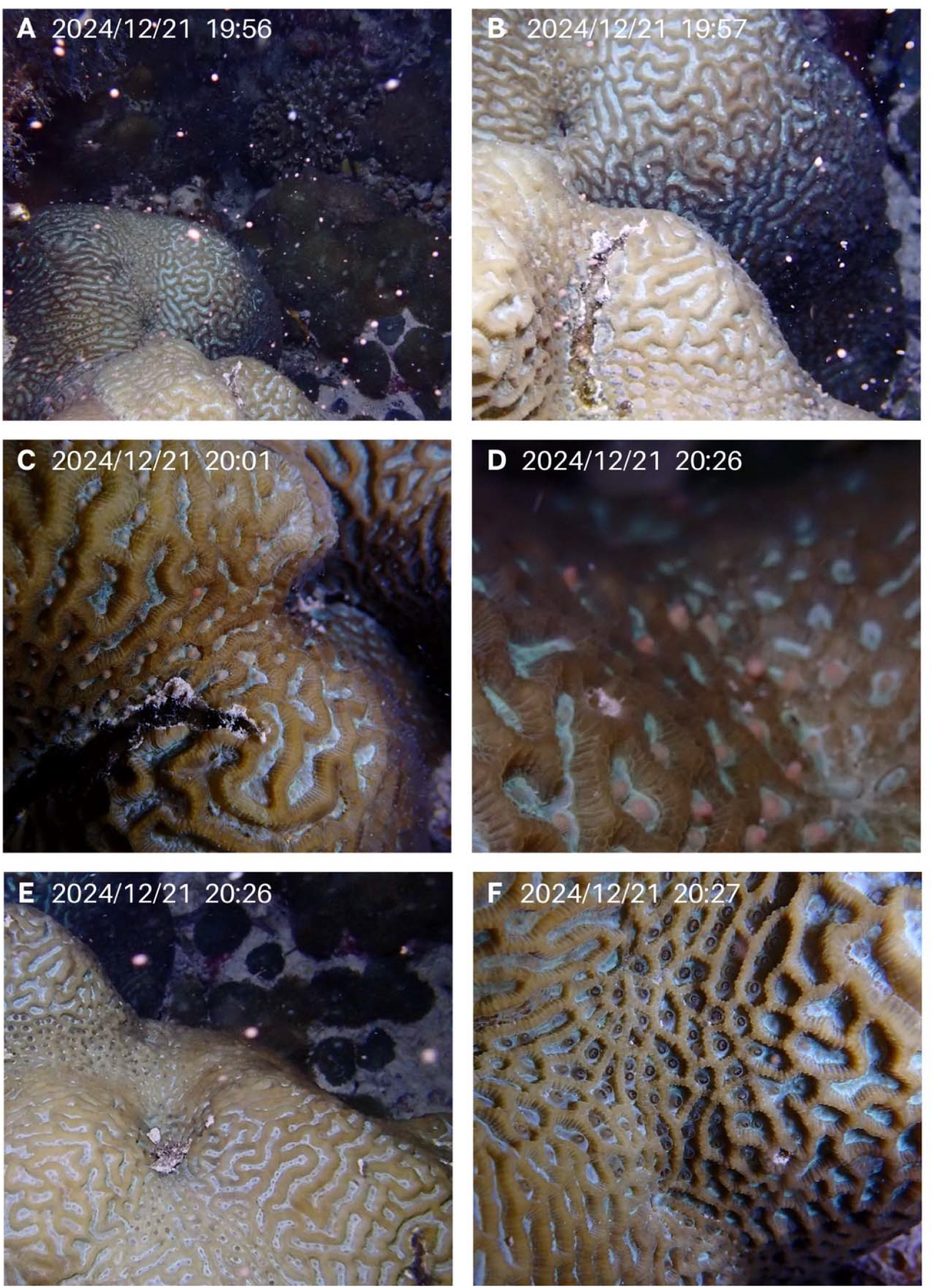
*Platygyra daedalea* spawning recorded in December 2024. A and B show initial rapid release of gamete bundle cloud. Gamete setting is seen in C and D before the second rapid release observed E. Panel F shows empty gaping polyp mouths following the release.

**Figure 5.**
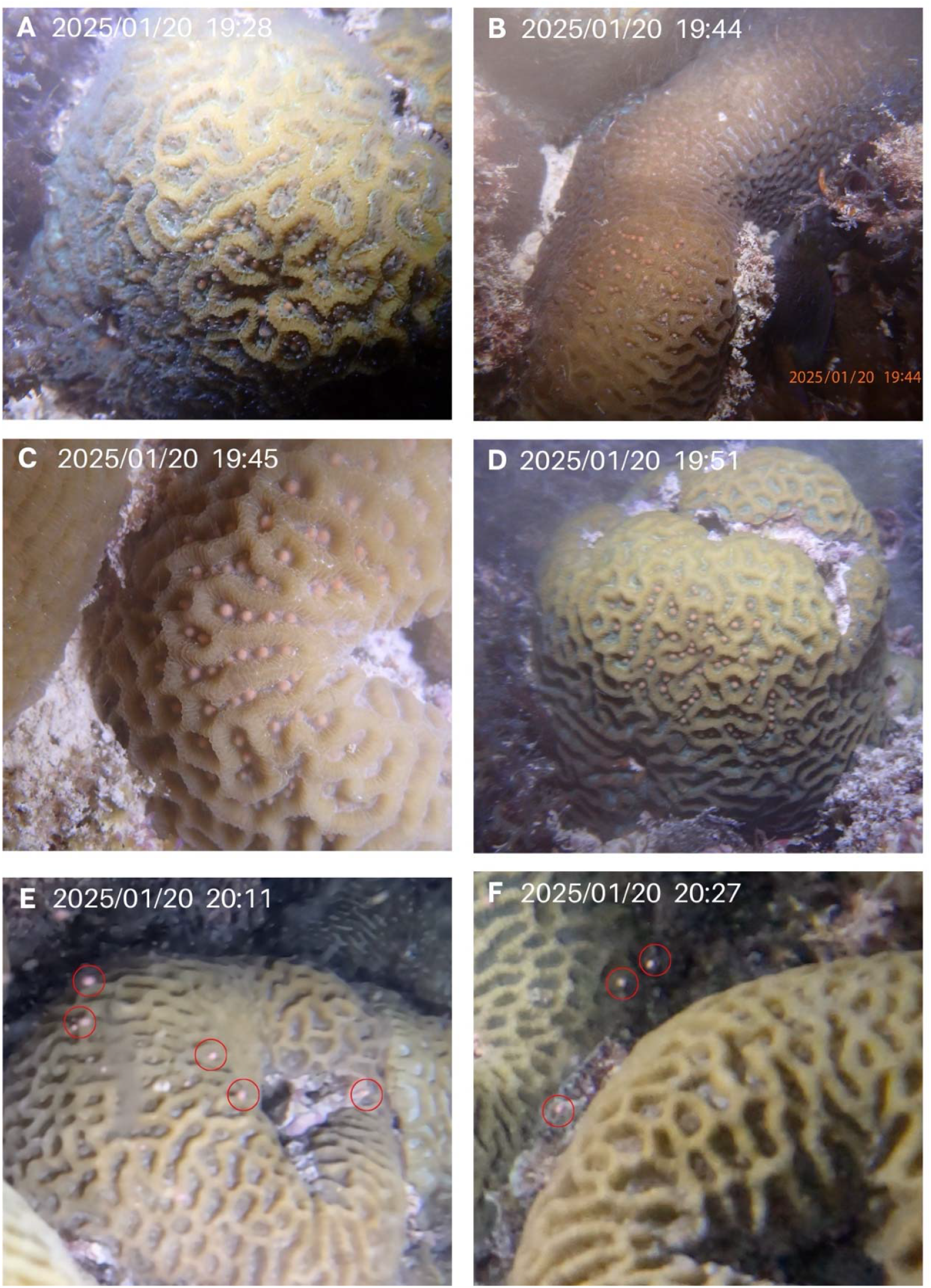
*Platygyra daedalea* spawning sequence in January 2025, observed in several colonies. Red circles indicate gamete bundles in the water column. A-D show gamete setting for four colonies, gamete bundles are released in E and F.

One individual colony was observed spawning in December, while four were documented to spawn in January, including the individual that spawned the previous month (Fig. 5). In January, a small colony showed visible pink gamete bundles setting starting at 19:28 with other adjacent colonies exhibiting setting gamete bundles within the next 15 minutes. Setting lasted 38-40 minutes, and gamete bundles were rapidly released by each colony in less than 2 minutes (Fig. 5 D,E).

## 4. Discussion

This study presents new *in situ* documentation of spawning timing and behaviour for *G. astreata, F. pentagona*, and *P. daedalea* in the Western Indian Ocean, specifically coastal Zanzibar. These findings provide a valuable baseline for coral scientists, conservationists, and restoration practitioners, and support future monitoring efforts in the region. Though limited to one site over four months, our observations contribute novel insights into coral spawning in a region where such data have been scarce. They expand the known geographic range of spawning for these species, previously reported mainly in the Indo-Pacific, particularly Asia. For instance, *F. pentagona* and *P. daedalea* have been recorded spawning in Thailand, Singapore, and Taiwan (Baird *et al*. 2021, Lin and Nozawa, 2017), and *G. astreata* in Singapore (Baird *et al*., 2021). Our results partially align with these records but also reveal differences likely driven by local environmental conditions.

In Zanzibar, *G. astreata* spawned on the fourth night after the full moon in November and December, within a consistent time window (46–54 minutes after sunset). Presenting similar seasonality, with reproductive peaks in the time of most rapid temperature increases, Singapore records show spawning in April, 3–5 nights after the full moon (Baird *et al*. 2021), but with greater variation in timing. In 2012 and 2014, the Singapore-documented spawning by Tun and Low (unpublished data, Baird *et al*. 2021) comprised 10 *G. astreata* individuals, with the spawning time ranging from 47 to 157 minutes after sunset (median = 103). While only one *G. astreata* individual was observed in Zanzibar, the consistent timing we observed suggests intra-individual regularity but highlights the need for broader sampling to assess inter-individual variation. *F. pentagona* was also represented in Zanzibar by a single colony, which spawned in November on the sixth night after the full moon, 63 minutes after sunset. No subsequent spawning was observed December through February. In Taiwan, where the only other observations of *F. pentagona* exist, this species has been documented spawning only twice, in April 2019 on the fourth and sixth nights after the full moon, several hours after sunset (Baird *et al*. 2021). These limited data and the variability in spawning nights and times observed between Zanzibar and Taiwan in these species for which we have limited data underscore the need for further monitoring to assess population-level synchrony.

*P. daedalea* spawning was observed in multiple individuals, offering insights into intra-population synchrony at our site. A single colony spawned in December, while four of six colonies observed spawned in January—suggesting a peak in that month. No spawning was seen in February. All spawning occurred on the sixth night after the full moon, beginning around 80 minutes after sunset, with gamete release lasting just 1–3 minutes. This brief release window has implications for monitoring and gamete collection. In contrast, *P. daedalea* previously documented spawning in Taiwan, Thailand, and Singapore spawned 4-8 days after the full moon, at variable times past sunset, and none of the releases were recorded as rapidly occurring (Baird *et al*. 2021, Lin and Nozawa, 2017). Interestingly, Kenyan populations show peak spawning in February–March with a secondary peak in August–October (Mangubhai and Harrison, 2008), based on patterns in gametogenesis and not direct observations of spawning. These variations in seasonal timing might reflect spatial or temporal differences in environmental cues such as temperature, photoperiod, and tidal cycles, all of which can influence gametogenesis and spawning readiness (Lin and Nozawa, 2023). Further observation over multiple years and expanded spatial coverage is needed to understand the factors influencing reproductive timing.

Our findings reveal distinct species-specific spawning windows and behaviours, underscoring the complexity and ecological significance of coral reproductive strategies at local scales. Variation in gamete setting, release duration, and evidence of split spawning in some colonies was observed. These behavioural nuances further emphasize the importance of localized monitoring, as within the same reef system, reproductive strategies can differ among individuals and species. The presence of split spawning—where not all colonies spawn simultaneously—may serve as a bet-hedging strategy, increasing the likelihood of successful fertilization under variable conditions. However, it also presents challenges for coral restoration practices that rely on synchronous gamete collection.

By establishing baseline reproductive data for corals in Zanzibar, this study supports conservation strategies in a region facing growing environmental pressures. These spawning data can inform the timing of conservation and restoration interventions— such as closed areas, larval propagation, or managed breeding—ensuring they are aligned with natural reproductive windows to maximize success. Understanding this variability is vital for effective reef restoration and management, especially as reproductive timing may be sensitive to climate change. While this study provides an important first step, long-term, multi-site monitoring across more species is essential for a comprehensive understanding of reproductive dynamics in the WIO.

## 5. Acknowledgments

We express gratitude for the fieldwork support provided by the Zanzibar marinecultures.org team, without whom these observations would not have been possible. Thanks to marinecultures.org leadership, Christian Vaterlaus, Connie Sacchi, Thomas Sacchi, Fabienne Addor, and Ali Mahmudi Ali for their commitment to this project. We are especially grateful to the marinecultures.org night dive team members: Ali Pandu Suleiman (Tabu), Abdi Mjaka Haji, Haji Mjaka (Shamte), Mohamed Juma Haji (Hijabu), and Captain Hassan.

